# The essential inter-domain interaction between NUDT9H and channel domain of human TRPM2 is also accomplished in trans

**DOI:** 10.1101/2023.06.21.545868

**Authors:** Wiebke Ehrlich, Frank J.P. Kühn

**Author notes:** Corresponding author: Dr. Frank Kühn;.

## Abstract

Channel function of human transient receptor potential melastatin type 2 (hsTRPM2) essentially depends on its C-terminal domain NUDT9H, which is homologous to the human Nudix hydrolase NUDT9. This cytosolic enzyme specifically binds and cleaves adenosine 5′-diphosphate ribose (ADPR), which in turn represents the principal agonist of TRPM2. For hsTRPM2 the experimental data strongly suggest, that binding of ADPR to NUDT9H as well as to a separate N-terminal binding pocket induces channel gating. Recent cryogenic electron microscopy (cryo-EM) analyses have provided the first concrete clues as to how NUDT9H interacts with the channel domain. In the present study we take an alternative approach by testing co-expression of NUDT9H together with a C-terminally truncated non-functional variant of hsTRPM2. Our data obtained from co-immunoprecipitation and proximity ligation assays reveal that NUDT9H and channel domain also specifically interact when co-expressed as independent proteins. Most importantly, calcium imaging as well as whole-cell patch-clamp recordings demonstrate that this in-trans interaction restores channel function, after stimulation either with intracellular ADPR or with extracellular hydrogen peroxide. Moreover, point mutation N1326D within the NUDT9H domain previously shown to be essential for TRPM2 function significantly reduces co-immunoprecipitation of NUDT9H as well as ADPR-dependent channel activity. These findings open up new possibilities to identify the molecular determinants of this crucial inter-domain interaction.

## Introduction

TRPM2 belongs to the melastatin subgroup of transient receptor potential (TRP) channels which are classified to the large family of cation-selective ion channels. TRP channels are characterized by a wealth of unusual properties, which are particularly reflected in their diverse activation mechanisms. In addition, these channels are essentially involved in numerous physiological processes (reviewed e. g. in Koivisto et al., 2022). A common theme seen in biology is that structure determines function which in a special way also applies to hsTRPM2. This ion channel plays a key role in the process of oxidative stress mediated apoptosis (e. g. reviewed in Malko et al., 2021). For this purpose, the TRPM2 channel protein shows a unique structure where the C-terminus contains a domain which is derived from the human Nudix hydrolase NUDT9 (Perraud et al., 2001). NUDT9 represents an ADPR pyrophosphatase which specifically cleaves the nicotinamide adenine dinucleotide (NAD^+^) metabolite ADPR in adenosine monophosphate and D-ribose 5’phosphate (Lin et al., 2002; Perraud et al., 2003; Shen et al., 2003).

It has been shown that in human cells oxidative stress induces poly-ADP-ribosylation of DNA and various nuclear proteins which is catalyzed by poly-ADPR polymerase type 1 (PARP-1). In the further course of this process the attached poly-ADPR chains are degraded to monomeric ADPR by poly-ADPR-glycohydrolase (PARG; reviewed e. g. in Pascal and Ellenberger, 2015). The resulting increase in the cytosolic concentration of ADPR in turn activates hsTRPM2 because ADPR represents the principal agonist of this non-selective plasma membrane cation channel (Perraud et al., 2001, 2005). The stimulation of hsTRPM2 in particular causes a massive influx of calcium through the channel pore ultimately leading to cell death (Fonfria et al., 2004; Miller, 2004). This indirect activation mechanism by oxidative stress can be experimentally simulated by extracellular application of H_2_O_2_ (Hara et al., 2002).

For a long time, the detailed gating mechanism of hsTRPM2 induced by ADPR was a mystery. Significant progress was recently made after functional analysis of far distantly related species variants of hsTRPM2 (Kühn et al., 2015, 2016, 2019a, 2019b; Iordanov et al., 2019; Tóth et al., 2020) together with cryo-EM studies (Huang et al., 2018, 2019; Wang et al., 2018; Yu et al., 2021; Huang et al., 2023). The resulting data reveal the presence of two distinct ADPR binding pockets, one in the NUDT9 homology domain (NUDT9H) and one in the N-terminal part of the channel. In TRPM2 channels of cnidaria and choanoflagellates NUDT9H retains full ADPRase activity but is not essential for ADPR-dependent channel gating, whereas in TRPM2 of human and zebrafish NUDT9H is catalytically inactive but indispensable for channel activation (reviewed by Kühn, 2020; Szollosi, 2021). Apparently, the interaction between channel domain and NUDT9H has changed significantly during metazoan evolution (Kühn et al., 2017). In archaic TRPM2 channels, the NUDT9H domain both is involved in pore opening and regulation of channel activity (Huang et al., 2023). However, the regulatory role seems to predominate here, since these channels work as well even without NUDT9H domain (Kühn et al., 2016; Iordanov et al., 2019; Huang et al., 2023). In contrast, in advanced species variants of TRPM2 the catalytic activity and thus the regulatory role of NUDT9H has been lost, whereas the structural interaction between NUDT9H and channel domain was crucially expanded (Huang et al., 2019; Huang et al., 2023).

Notwithstanding recent major advances in elucidating the gating mechanism of TRPM2, a number of unanswered questions remain. For example, there is ambivalence regarding the relative importance of both ADPR binding sites for channel activation as well as about the substrate specificities among different species variants (reviewed by Kühn, 2020). In particular, for hsTRPM2 it remains to be determined which amino acid residues of NUDT9H accomplish the essential interaction with the channel domain. Although cryo-EM studies have identified certain subdomains of full-length hsTRPM2 that are involved in this interaction (Wang et al., 2018; Huang et al., 2019) the functional verification of all the postulated molecular determinants is still pending. In addition, a new study strongly suggests that in hsTRPM2 especially the Cap-region of NUDT9H is crucial (Huang et al., 2023). So far, only the point mutation N1326D has been described which is localized within the Cap region of NUDT9H, outside of the actual ADPR binding site preventing ADPR-dependent channel activation of hsTRPM2 (Kühn and Lückhoff, 2004; Du et al., 2009).

To be able to analyze the effects of point mutations on structural and functional interactions side by side, we tested the possibility of expressing both domains separately and allowing them to interact in trans. With this novel approach, we aim to provide a simplified way to further advance the identification of crucial amino acid residues involved in the interaction between NUDT9H and channel domain. With two different detection methods we can demonstrate that in hsTRPM2 channel domain and NUDT9H also specifically interact when co-expressed as independent proteins. Most importantly, the separate NUDT9H protein largely restores channel activity of the truncated and non-functional channel fragment, as revealed by calcium imaging and whole-cell patch clamp analysis. Exemplified by the modification N1326D, we were able to show that this mutation actually impairs both the structural and the functional interaction between NUDT9H and channel domain. These results should pave the way for further investigations in order to precisely characterize the functional cooperation between these two domains in hsTRPM2.

## Experimental procedures

### Molecular Biology

For interaction studies of C-terminally truncated TRPM2 channel with different NUDT9H variants as well as with NUDT5 as a negative control, the following constructs were generated. The cDNA of hsTRPM2-ΔNUDT9H-3xHA (EGFP) subcloned as EcoRI-XbaI fragment into pIREShrGFP-2a vector (Agilent, Santa Clara, CA, USA) was prepared as described by Kühn et al. 2016. For generation of hsNUDT9H-FLAG (DsRed) the pIREShrGFP-2a vector containing hsTRPM2-1xHA (Kühn et al., 2016) was used and the single hemagglutinin (HA)-tag was changed into a single FLAG-tag by two separate mutagenesis steps (step I: 5’-aagtacccatatgacgtt-3’ to 5’-gattacaaggatgaccat-3’; step II: 5’-ccagactac-3’ to 5’-gataagtag-3’). Then, a second EcoRI restriction site was introduced in front of the open reading frame of hsNUDT9H including a N-terminal linker sequence of about 70 amino acid residues. Subsequently, the DNA sequence of hsTRPM2 upstream of the reconstructed open reading frame of hsNUDT9H (starting at methionine 1173 ending with tyrosine 1503) was deleted via EcoRI double restriction digest and the DNA fragment containing the sequence of vector and NUDT9H-FLAG was re-ligated. In a final step EGFP was replaced by DsRed. Therefore, the open reading frame of EGFP was deleted from the vector by double digest with HindIII + ScaI and replaced by the corresponding HindIII + ScaI DNA fragment containing the open reading frame of DsRed. The open reading frame of human NUDT5 (Gene banc Acc. No. CAG33476.1) flanked with appropriate 5’ and 3’ restriction sites (5’-NruI and 3’-XbaI) was synthesized by Eurofins scientific (Ebersberg, Germany) and subcloned into pIREShrGFP-2a-hsNUDT9H-FLAG (DsRed), after the open reading frame of hsNUDT9H has been deleted by digestion with NruI + XbaI. For generation of the channel variant hsNUDT9H-N1326D, an asparagine at position 1326 of hsNUDT9H-FLAG (DsRed) was replaced with aspartate using QuikChange mutagenesis system (Agilent). Defined oligonucleotides in sense and antisense configuration were obtained from (Eurofins). Each point mutation or cloning step was verified by DNA sequencing (Eurofins).

### Cell culture and transfection

For testing the specific in-trans interaction between NUDT9H and channel domain of hsTRPM2 both proteins were expressed in HEK-293 cells, either separately (negative control) or in combination and analyzed by the investigation methods described below. As a further negative control in some experiments NUDT9H was replaced by NUDT5.

Human embryonic kidney (HEK-293) cells were purchased from the German Collection of Microorganisms and Cell Cultures (Braunschweig, Germany). Cell culture was carried out in DMEM media (Biochrome, Berlin, Germany) supplemented with 4 mM L-glutamine, 2 mM sodium pyruvate and 10% (v/v) foetal calf serum (Biochrome). Transfection of HEK-293 cells was performed with FuGene 6 transfection reagent (Roche, Mannheim, Germany) according to the manufacturer’s protocol. To achieve an optimal co-expression the volumes of FuGENE 6, FCS-free culture medium and amount of DNA were adapted in an appropriate manner as summarized in **Tabs. 1–3**. Transfected cells were incubated for 24 h at 37°C in a 5% CO_2_ atmosphere. Subsequently, the cells were harvested for co-immunoprecipitation (Co-IP)experiments. Alternatively, the cells were seeded on poly-lysine-coated glass coverslips at a suitable dilution and incubated for 3–4 h. Then, whole-cell patch clamp recordings or calcium imaging experiments were performed with cells visibly positive for EGFP-and/or DsRed-expression. For protein-protein interaction studies the expressed proteins contain C-terminally attached tags (3xHA or FLAG) for immuno-detection, whereas for functional studies the corresponding tags were omitted. For proximity ligation assay the expression of EGFP and DsRed was abolished by introduction of premature stop codons into the corresponding open reading frames. An overview of the variants used is summarized in **Tabs. 1–3**. At least three independent transfections were used for each experimental group.

**Table 1:** Amount of DNA, FuGENE 6 and FCS-free culture medium used for transfection of HEK-293 cells for Co-Immunoprecipitation experiments.

| DNA channel core domain [μg] |  | DNA NUDT9H domain [μg] |  | FuGENE 6 [μl] | FCS-free culture medium [μl] |
| --- | --- | --- | --- | --- | --- |
| hsTRPM2ΔNUD-3xHA (green) | 7.5 | - |  | 20 | 428 |
| - |  | hsNUD-FLAG (red) | 5.6 | 15 | 319 |
| hsTRPM2ΔNUD-3xHA (green) | 7.5 | hsNUD-FLAG (red) | 5.6 | 35 | 747 |
| - |  | hsNUD-N1326D-FLAG (red) | 5.6 | 15 | 319 |
| hsTRPM2ΔNUD-3xHA (green) | 7.5 | hsNUD-N1326D-FLAG (red) | 5.6 | 35 | 747 |
| - |  | NUDT5-FLAG (red) | 6.5 | 15 | 319 |
| hsTRPM2ΔNUD-3xHA (green) | 7.5 | NUDT5-FLAG (red) | 6.5 | 35 | 747 |

**Table 2:** Amount of DNA, FuGENE 6 and FCS-free culture medium used for transfection of HEK-293 cells for Calcium Imaging and whole-cell patch-clamp experiments.

| DNA channel core domain [μg] |  | DNA NUDT9H domain [μg] |  | FuGENE 6 [μl] | FCS-free culture medium [μl] |
| --- | --- | --- | --- | --- | --- |
| hsTRPM2 (green) | 10 | - |  | 27 | 428 |
| hsTRPM2ΔNUD (green) | 10 | - |  | 27 | 570 |
| hsTRPM2ΔNUD (green) | 7.5 | hsNUD (red) | 5.6 | 35 | 747 |
| hsTRPM2ΔNUD (green) | 7.5 | hsNUD-N1326D (red) | 5.6 | 35 | 747 |

**Table 3:**
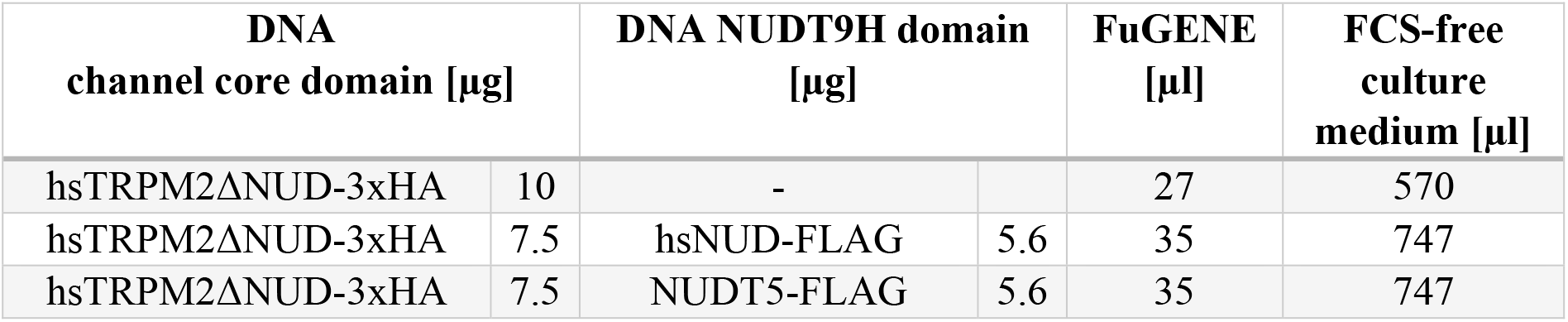
Amount of DNA, FuGENE 6 and FCS-free culture medium used for transfection of HEK-293 cells for proximity ligation assay.

| DNA channel core domain [μg] |  | DNA NUDT9H domain [μg] |  | FuGENE [μl] | FCS-free culture medium [μl] |
| --- | --- | --- | --- | --- | --- |
| hsTRPM2ΔNUD-3xHA | 10 | - |  | 27 | 570 |
| hsTRPM2ΔNUD-3xHA | 7.5 | hsNUD-FLAG | 5.6 | 35 | 747 |
| hsTRPM2ΔNUD-3xHA | 7.5 | NUDT5-FLAG | 5.6 | 35 | 747 |

### Coating of coverslips for fixation of HEK-cells

For whole-cell patch clamp or calcium imaging experiments, transfected HEK-cells were seeded on polylysine-coated coverslips in an appropriate dilution. For this purpose, the cover slips were soaked in methanol and then flame-scarfed. Subsequently, the cleaned coverslips were transferred into 34 x 10 mm culture dishes and covered with 50 µl polylysine (0.1 mg / ml, diluted in H_2_O). After 60 min of incubation at room temperature the excess polylysine was removed by two washing steps in PBS, followed by the addition of the cell suspension.

### Co-Immunoprecipitation

The Pierce™ HA-Tag IP/Co-IP Kit (Thermo Fisher Scientific, USA) was used according to the manufacturer’s instructions but with minor modifications. A single 100 x 20 mm culture dish of transfected, sub-confluent (80%) HEK-293 cells was used per test run. In brief, cells were lysed in CelLytic M (Sigma Aldrich, USA) lysis buffer containing 1 % protease inhibitor (Sigma Aldrich, USA). After incubation for 15 min on ice on a shaker, cells were removed from culture dish using a cell scraper. Cell suspension was centrifuged for 15 min at 12.000 x g at 4°C and the clarified supernatant was transferred to new tubes. Protein concentration was determined using Bradford reagent (Sigma Aldrich, USA) according to manufacturer’s instructions. 600 µg of sample protein were incubated with anti-HA agarose and after several washing steps elution was performed with SDS sample buffer (Thermo Fisher Scientific, USA). 25 µl each of the eluate samples and 9 µg of the corresponding cell lysates, used as input control, were subjected to SDS-PAGE (Bolt 4-12% Bis-Tris Plus Gel, Invitrogen) and Western blot analysis as previously described (Kühn et al., 2019b). Sometimes the samples were stored overnight at 4°C and analyzed the next day by SDS-PAGE and Western blot analysis.

PVDF Membranes were cut in halves between the marker bands for 50 and 65 kDa. The upper part of the membrane was incubated with a primary monoclonal mouse-anti-HA antibody (1:1000; H3663; Sigma-Aldrich, USA) to detect HA-tagged TRPM2 channel domain and the lower part of the membrane was incubated with rabbit-anti-FLAG (DYKDDDDK) antibody (1:1000; 701629; Invitrogen, USA) to detect FLAG-tagged NUDT9H domain and NUDT5, respectively. Both membrane parts were incubated with a rabbit-anti-mouse-(1:1000; P0261; DAKO, Agilent, USA) and swine-anti-rabbit-HRP conjugated secondary antibody (1:1000; P0217; DAKO, Agilent, USA). Detection of antibodies was performed using Intas Infinity ECL Starlight (Intas, Germany) and ChemoStar Touch (Intas, Germany) imaging system, for signal quantification ImageJ was used.

### Proximity Ligation Assay

For proximity ligation assay transfected HEK-293 cells were seeded in an appropriate dilution on Nunc Chamber Slides (Thermo Fisher, USA). After fixation in 4% paraformaldehyde for 15 min at RT, cells were washed 3 times for 5 min in TBS-T (TBS + 0.1 % Tween20). Subsequently, cells were permeabilized in PBS + 0.1 % Triton x-100 for 10 min at RT and then washed 3 times for 5 min in TBS-T at RT. The Duolink^®^ In Situ Orange Starter Kit mouse/rabbit (Sigma Aldrich, USA) was used according to the manufacturer’s instructions. Fluorescence signals were examined by fluorescence microscopy using a Zeiss LSM 700 microscope.

### Confocal microscopy of transfected HEK cells

Transfected HEK-cells were seeded on poly-L-lysin coated glass coverslips. After incubation for 3.5 h at 37 °C the cells were washed three times with PBS and then fixed for 45 min in 4 % paraformaldehyde at RT. Cells were then washed three times with PBS and air dried at RT. Afterwards cell-loaded coverslips were mounted. All steps were carried out in the dark. Fluorescence signals were examined by fluorescence microscopy using a Zeiss LSM 700 microscope.

### Calcium imaging

For fluorescence imaging of cytosolic calcium hereafter abbreviated to [Ca^2+^]_i_, transfected HEK-293 cells plated on poly-lysine-coated glass coverslips were loaded for 20 min at 37 °C in standard bath solution containing membrane-permeable Fura-2 acetoxymethylester (1.5 ng/µl; Invitrogen) and pluronic acid (0,025%). Fluorescence was alternately excited at 340 and 380 nm using the Polychrome IV monochromator (TILL Photonics). The emitted fluorescence was measured at 510 nm using a Sensicam (IMAGO). Fluorescence was corrected for background at each wavelength. Measurements were obtained at room temperature (21 °C). Activation of TRPM2 was elicited either by adding 1 µl of H_2_O_2_ (30% stock solution) directly to the bath solution (dilution about 1:1000) or by exchanging the bath with a standard bath solution already supplemented with 10 mM H_2_O_2_. Both forms of treatment had the same effect.

### Electrophysiology

Whole-cell patch clamp recordings of transfected HEK-293 cells were performed using an EPC 9 amplifier equipped with a personal computer with Pulse 8.5 and X Chart software (HEKA, Lamprecht, Germany). The standard bath solution contained (in mM) 140 NaCl, 1.2 MgCl_2_, 1.2 CaCl_2_, 5 KCl, 10 HEPES, pH 7.4 (NaOH). For Na^+^ free solutions, Na^+^ was replaced by 150 mM N-methyl-D-glucamine (NMDG) and the titration was performed with HCl. The pipette solution contained (in mM) 145 CsCl, 8 NaCl, 2 MgCl_2_, 10 HEPES, pH 7.2 (CsOH) and the Ca^2+^-concentration was adjusted to 1 µM (0.886 mM Ca^2+^, 1 mM Cs-EGTA). For the stimulation of TRPM2, adenosine 5’-diphosphate ribose (ADPR, Sigma-Aldrich, Munich, Germany) was prepared as 100 mM stock solutions in distilled water and aliquots stored at – 20°C. On the day of the experiment the ADPR stock solution was diluted to the desired concentration in the intracellular (pipette) solution. Stimulation wit H_2_O_2_ (10 mM, diluted from a 30% stock solution) was carried out in the same way as described in calcium imaging experiments (see above). In this case the pipette solution contains no ADPR. Unless otherwise stated, the experiments were performed at room temperature (21°C) and the current-voltage relations were obtained during voltage ramps from –150 to +150 mV and back to –150 mV applied over 200 ms. The holding potential was –60 mV. For the analysis the maximal current amplitudes (pA) in a cell were divided by the cell capacitance (pF), as measure of the cell surface, resulting in the current density (pA/pF).

### Data analysis and statistics

The comparison of two groups in calcium imaging experiments and whole-cell patch clamp experiments was performed using a two-tailed Mann Whitney Test and the data are expressed as the mean ± SEM. Differences were considered significant at **P < 0.01, ***P < 0.005 and ****P < 0.0001. In Co-IP experiments an unpaired Student’s t-test with Welch’s correction was performed and the data are expressed as the mean ± SM. Differences were considered significant at **P <0.01, ****P < 0.0001.

## Results

### Co-expression of channel domain and NUDT9H domain as separate proteins

A sequence comparison between the human NUDT9 enzyme (hsNUDT9) and the NUDT9 homology (NUDT9H) domains of the TRPM2 species variants of human, zebrafish and sea anemone revealed that in the respective homology domains a stretch of about 60 amino acid residues of the N-terminus of hsNUDT9 is almost completely absent (Kühn et al., 2017). On the other hand, a sequence comparison with hsTRPM8, the most closely related TRP channel of hsTRPM2 (Peier et al., 2002), which does not contain a NUDT9H domain, strongly suggests that in hsTRPM2 there is a linker of approximately 70 amino acid residues in length between channel domain and NUDT9H domain which is hardly homologous to hsNUDT9 (Kühn et al., 2010). Despite the low homology, we decided to assign this linker region to the actual NUDT9H domain. Therefore, we included this sequence into the open reading frame of the separately expressed NUDT9H domain of hsTRPM2 using a naturally occurring methionine residue (M1173) in the outermost N-terminal region of this linker as start codon (ref. to Experimental procedures).

For the protein-protein interaction studies as well as the functional tests, the following proteins were heterologously expressed in HEK-293 cells, either separately or in pairs of two: (1) The truncated hsTRPM2 channel without NUDT9H domain, optionally equipped with a triple HA-tag attached to the C-terminus. Positive expression was monitored by green fluorescence (EGFP-expression; **Fig. 1, left**). (2) The separate NUDT9H domain of hsTRPM2, optionally with a FLAG-tag C-terminally attached. Positive expression was monitored by red fluorescence (DsRed-expression; **Fig. 1, middle**). As a further variant of this protein a mutant was used containing the point mutation N1326D. (3) The human Nudix hydrolase 5 (hsNUDT5), a paralogue of NUDT9H which is not known to interact with any ion channel (used as negative control). This protein was optionally FLAG-tagged and was also monitored by red fluorescence (DsRed-expression). Representative confocal microscopy images of co-transfected HEK-293 cells are depicted in **Fig. 1**. The pictures demonstrate that the transfection level was consistently high, as was to be expected with a transfection system that has been established in our laboratory for many years. A high expression level was particularly desirable for co-expression experiments in order to obtain clear signals both in functional tests and in protein-protein interaction studies. For this purpose, the transfection protocol was optimized for each experimental set up **(Tabs. 1-3)**. Successful co-expression was indicated by the relative number of yellow fluorescent cells, with a small number of orange-stained cells also being detected (**Fig. 1, right**). Especially for the functional tests, only cells with strong yellow fluorescence were analyzed.

**Figure 1.**
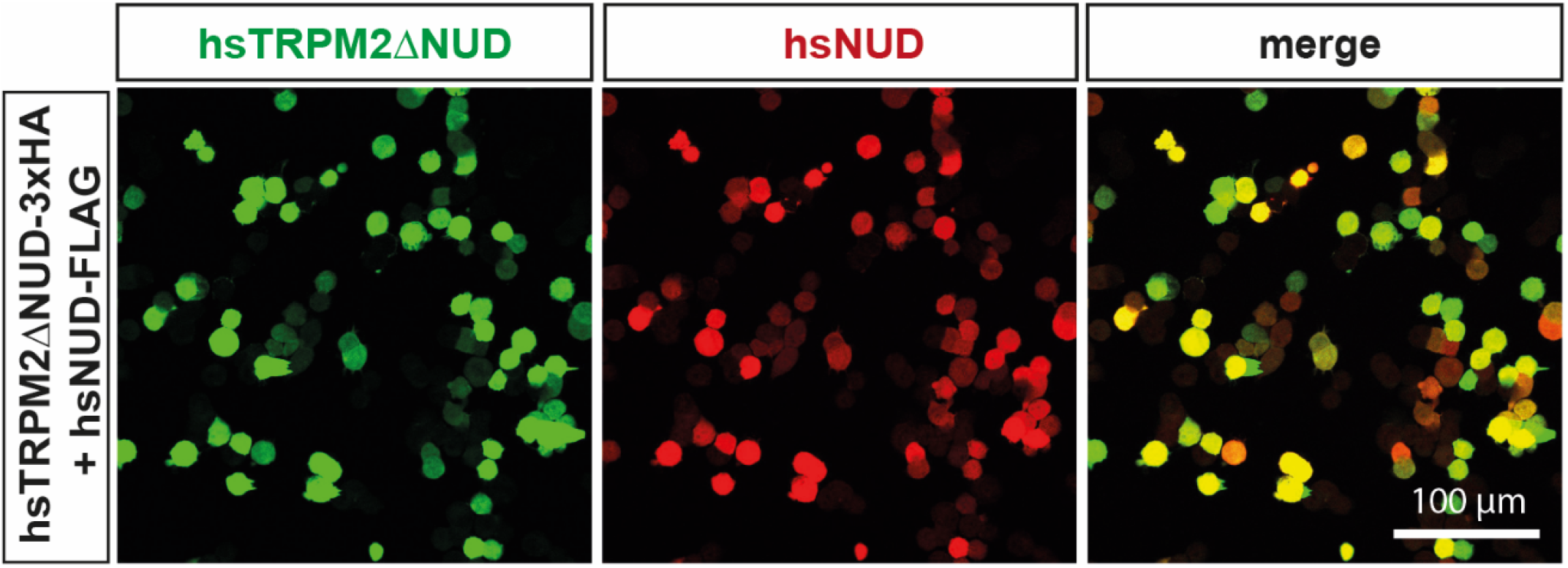
Representative confocal microscopy images of co-transfected HEK-293 cells obtained with different fluorescence filters. The successful expression of hsTRPM2ΔNUDT9H and hsNUDT9H is visualized by using specific fluorescence filters. Expression of hsTRPM2ΔNUDT9H **(left)** and hsNUDT9H **(middle)** is monitored by green (EGFP) or red (DsRed) fluorescence signals. The stronger the yellow fluorescence signal, the more extensive is the co-expression in the respective cell **(right)**. Especially for functional studies (calcium-imaging and whole-cell patch-clamp recordings) of co-expression only the yellow cells were analyzed. Images were taken at 40x magnification, the scale bar can be applied for all images

### Co-immunoprecipitation studies

To test for a specific protein-protein interaction of C-terminally truncated hsTRPM2 and hsNUDT9H we performed co-immunoprecipitation studies. The rational of this experiment was to immobilize the 3x-HA-tagged hsTRPM2-ΔNUDT9H channel domain with anti-HA agarose. In case of a strong interaction between co-expressed proteins (cell lysates of yellow cells) the FLAG-tagged putative interaction partner should be co-immobilized. After performing several stringent washing-steps the eluate fraction was submitted to SDS gel electrophoresis followed by Western-blot analysis with anti-HA or anti-FLAG antiserum. **Fig. 2** shows a representative Western-blot of our co-immunoprecipitation experiments including the corresponding control data. In the right panel of **Fig. 2A** the cell lysates of single-transfected or co-transfected HEK-293 cells were applied as indicated and screening was performed with anti-HA-antiserum (upper part) or with anti-FLAG-antiserum (lower part). The detected signals (input control) showed comparable levels of expression for each protein either when expressed separately or in combination, even if these levels were subjected to a certain fluctuation in repeated experiments The eluate fractions (**Fig. 2A**, left panel) showed strong signals of 3x-HA tagged hsTRPM2-ΔNUDT9H which indicates significant enrichment by anti-HA agarose treatment, if compared with the corresponding cell lysates. Separately expressed FLAG-tagged hsNUDT9H (wild-type or mutant) as well as FLAG-tagged hsNUDT5 were not detected in the eluate fraction indicating the absence of non-specific interactions between FLAG-tagged proteins and anti-HA-agarose. However, a robust signal for hsNUDT9H-FLAG was detectable in the eluate fraction when this protein was co-expressed with hsTRPM2-ΔNUDT9H-3xHA. Likewise, a positive signal was visible for the co-expressed variant of the hsNUDT9H domain containing point mutation N1326D, albeit significantly reduced compared to wild-type hsNUDT9H (**Fig. 2B**). As expected for co-expressed hsNUDT5 no signals could be detected in the eluate fraction with anti-FLAG antiserum indicating the absence of specific protein-protein interactions with 3x-HA tagged hsTRPM2-ΔNUDT9H (**Fig. 2A, B**).

**Figure 2.**
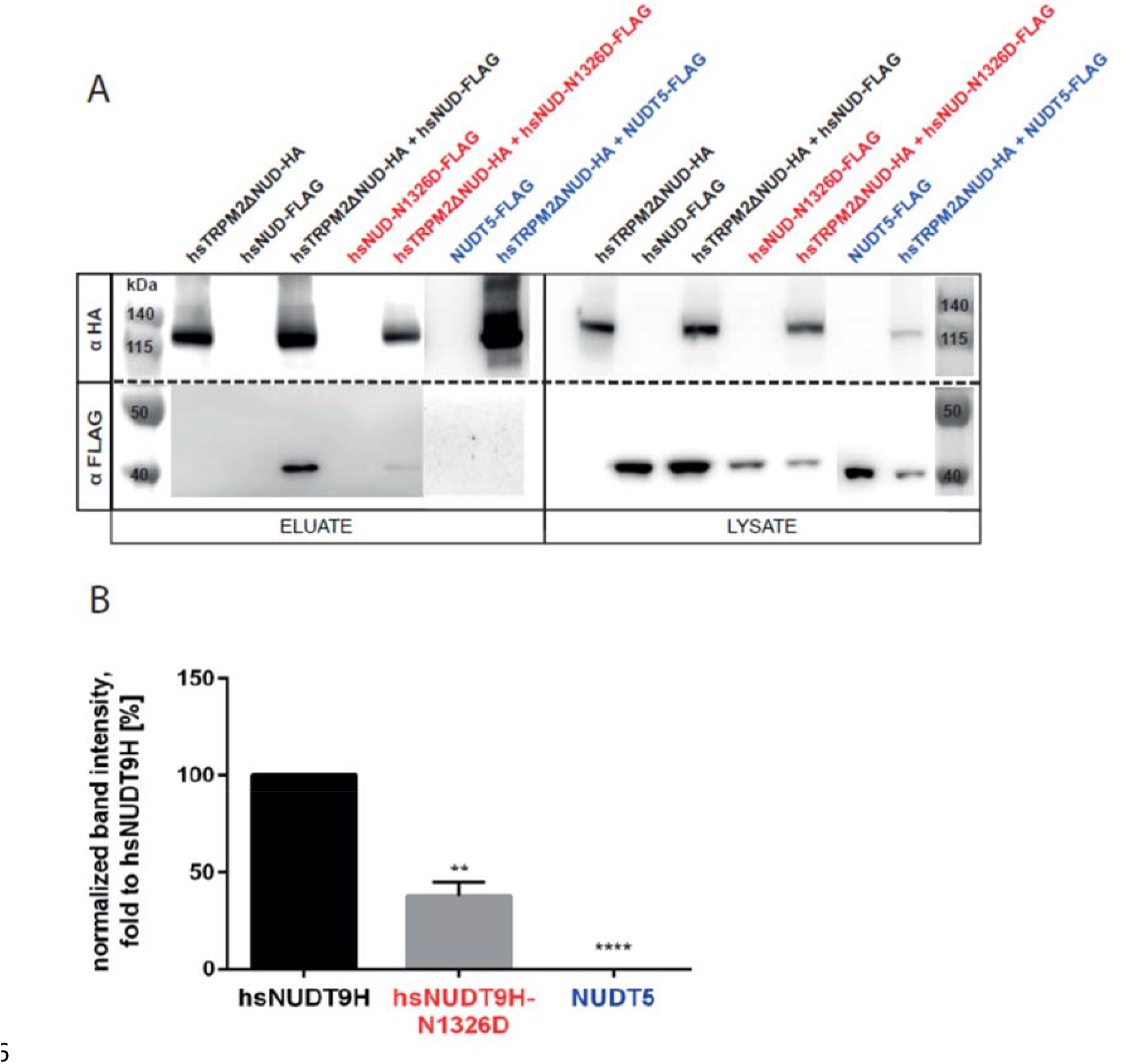
Representative data of Co-IP experiments with HEK-293 cell lysates expressing hsTRPM2ΔNUDT9H together with one of two variants of NUDT9H or together with NUDT5. HEK-293 either transfected with the individual proteins or in combinations (as indicated) were subjected to Co-IP with subsequent SDS-PAGE and Western blot. For Co-IP via anti-HA agarose as well as for immunohistochemically staining using anti-HA-or anti-FLAG antiserum the individual proteins were expressed with the corresponding C-terminal 3x-HA tags and FLAG-tags, respectively. Total cell lysates (LYSATE) were analyzed to verify correct expression of all proteins tested. (**panel A, right**) Eluate fractions (ELUATE) of the loaded anti-HA agarose show positive signals for hsTRPM2ΔNUDT9H-3xHA, as expected. However, for (FLAG-tagged) putative interaction partners positive signals were only detected after successful Co-IP with hsTRPM2ΔNUDT9H-3xHA, indicating robust in-trans interaction between co-expressed proteins (**panel A, left**). The blot membrane was cut between 50 and 65 kDA (indicated by a dotted line). The upper part was screened with anti-HA antiserum and the lower part with anti-FLAG antiserum. **(Panel B)** Summary of repeated Co-IP experiments performed as exemplary shown in Fig. 2 panel A. Signal intensities were quantified using ImageJ software. The signal intensities of individual co-immuno-precipitated samples were normalized to the corresponding input control (total cell lysate fraction) and compared to the respective data of Co-IP of hsTRPM2ΔNUDT9H-3xHA and hsNUDT9H-FLAG. Averaged values with standard deviation of the mean (SM) are shown and statistical analysis was performed with an unpaired Student’s t-test with Welch’s correction, n = 3-9. p-values <0.0001 ****, <0.01 **. Number of independent experiments (n): hsTRPM2ΔNUDT9H-3xHA + hsNUDT9H-FLAG, (n=9); hsTRPM2ΔNUDT9H-3xHA + hsNUDT9H-N1326D-FLAG (n=4); hsTRPM2ΔNUDT9H-3xHA + hsNUDT5-FLAG (n=3).

### Analysis of specific protein-protein interaction by proximity ligation assay (PLA)

The PLA method utilizes oligo-coupled antibodies as detection or proximity sensors for target antigens. Using this sophisticated method, we wanted to confirm our results obtained from the CoIP experiments which already had indicated a positive in-trans interaction between hsTRPM2ΔNUDT9H and hsNUDT9H. With PLA a positive signal will be only generated if the two potential interaction partners come very close together which in turn strongly suggests a direct interaction. As shown in **Fig. 3A** there are clearly positive signals for protein-protein interaction (red fluorescence) in HEK-293 cells co-expressing hsTRPM2ΔNUDT9H + hsNUDT9H which are absent in the negative controls (**Fig. 3B, 3C**). These positive signals are also completely missing in cells transfected with hsTRPM2ΔNUDT9H alone (**Fig. 3D**) as well as in the corresponding negative controls (**Fig. 3E, 3F**). To verify that the observed fluorescence signals were obtained by specific binding of the putative interaction partners as well as to estimate the level of staining background, two different negative controls were used: (i) the secondary antibody was exclusively applied without primary antibody (**Fig. 3B, 3E**) (ii) the primary antibody was replaced by an isotype control (**Fig. 3C, 3F**) that lack specificity to the target.

**Figure 3.**
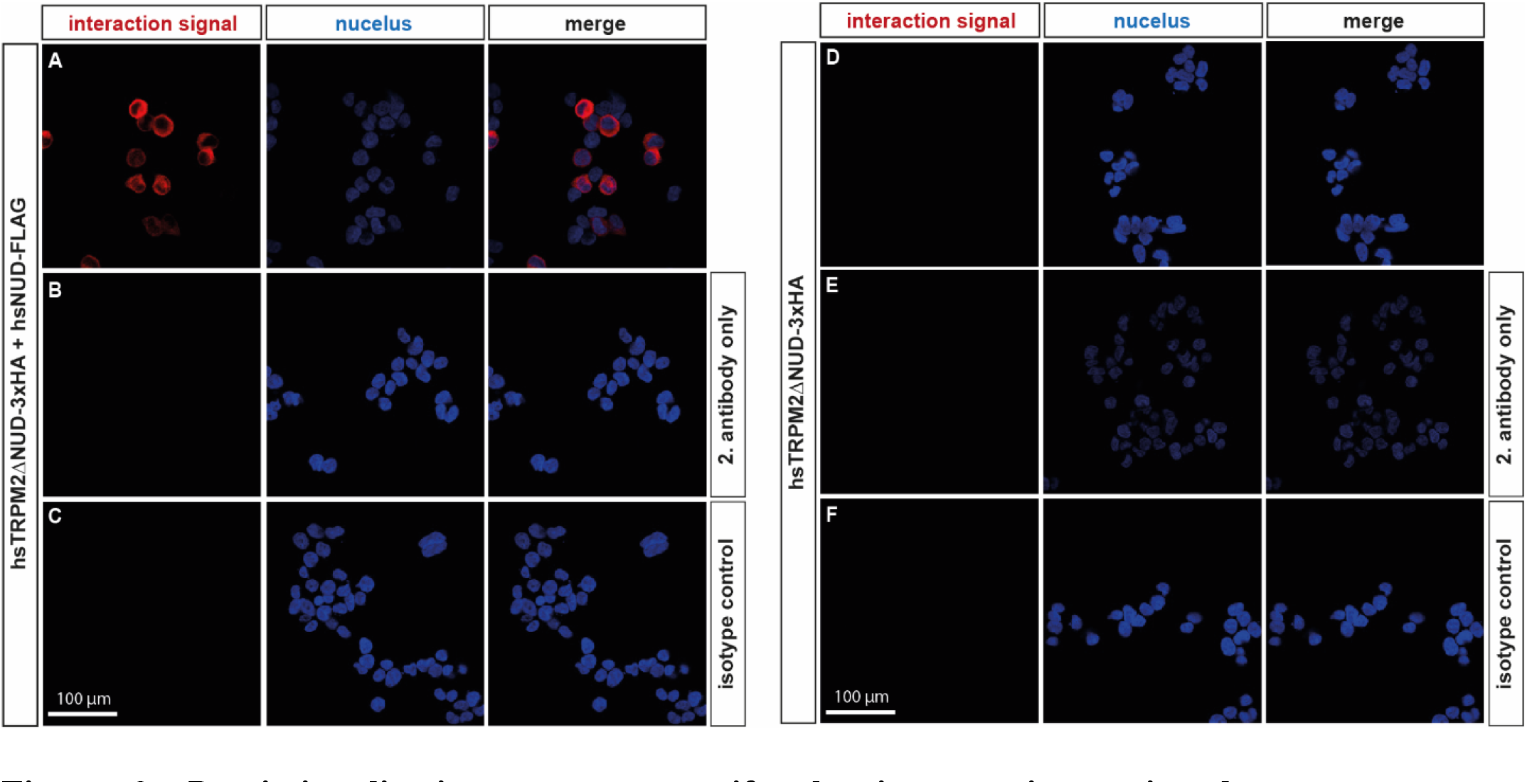
Proximity ligation assay to verify the in-trans interaction between hsTRPM2ΔNUDT9H-3xHA and hsNUDT9H-FLAG. Protein-protein interactions are indicated by red fluorescence signals which were exclusively detected in HEK-293 cells co-transfected with hsTRPM2ΔNUDT9H-3xHA + hsNUDT9H-FLAG **(panel A)**. In cells only expressing the C-terminal truncated channel hsTRPM2ΔNUDT9H-3xHA no positive signals for protein-protein interaction could be identified **(panel D).** Staining with the secondary antibody alone **(panels B and E)** or isotype controls (rabbit IgG/mouse IgG) **(panels C and F)** were used as negative controls. Images were taken with a Zeiss LSM 700 fluorescence microscope at 40x magnification, the scale bar can be applied for all images.

Thus, using two independent methods, we were able to demonstrate a specific in-trans interaction between NUDT9H domain and C-terminally truncated human TRPM2 channel. However, no indications can be derived from these data as to whether this interaction is also functionally relevant. This question was addressed in the subsequent experiments.

### Functional analysis of in-trans interaction by calcium imaging

For calcium imaging experiments HEK-293 cells were single-transfected with wild-type hsTRPM2 (positive control) and hsTRPM2ΔNUDT9H, respectively or co-transfected with hsTRPM2ΔNUDT9H + hsNUDT9H. As already described above, for all functional tests the corresponding proteins were expressed without C-terminal tags.

As a prerequisite for this experimental approach a stimulation from the extracellular side of the plasma membrane is mandatory. This is made possible by the fact that hsTRPM2 can be indirectly stimulated via extracellular application of H_2_O_2_. Since this kind of stimulation takes place via a gradual accumulation of intracellular ADPR (Perraud et al., 2005), it may be even more sensitive than the direct application of high concentrations of ADPR via the patch pipette during whole-cell patch clamp experiments. As a positive control, we used HEK-293 cells single-transfected with full-length wild-type hsTRPM2, which typically showed a consistent and long-lasting increase in [Ca^2+^]_i_ in response to H_2_O_2_ (**Fig. 4A**) as previously reported (e.g. Hecquet et al., 2007; Kühn et al., 2015). Similarly, in HEK-293 cells co-transfected with hsTRPM2ΔNUDT9H + hsNUDT9H, a considerable increase in [Ca^2+^]_i_ was detected after the application of 10 mM H_2_O_2_. This response was comparable to the positive control with respect to signal intensity and time course (**Fig. 4B**). In contrast, cells single-transfected with the C-terminal truncated variant of hsTRPM2 (hsTRPM2ΔNUDT9H) showed only a small increase in [Ca^2+^]_i_ in response to H_2_O_2_ (**Fig. 4C**) which was indistinguishable from that of mock-transfected cells (not shown). The results of several independent calcium imaging experiments are summarized in **Fig. 4D** indicating that co-expression of hsTRPM2ΔNUDT9H + hsNUDT9H largely restores channel function according to hsTRPM2. Since activation of TRPM2 essentially depends on the presence of extracellular Ca^2+^, the depletion of extracellular Ca^2+^ suppresses TRPM2-dependent Ca^2+^-influx into the cell (e.g. Kühn et al., 2010, 2015). As shown in **Fig. 5** the exchange of standard bath solution (1.2 mM Ca^2+^) to a divalent-free bath solution induced significant decrease of [Ca^2+^]_i_ both in cells single-transfected with hsTRPM2 (**Fig. 5A**) and co-transfected with hsTRPM2ΔNUDT9H + hsNUDT9H (**Fig. 5B**). In contrast, in cells single-transfected with hsTRPM2ΔNUDT9H the depletion of extracellular Ca^2+^ had no effect on [Ca^2+^]_i_ (**Fig. 5C**).

**Figure 4.**
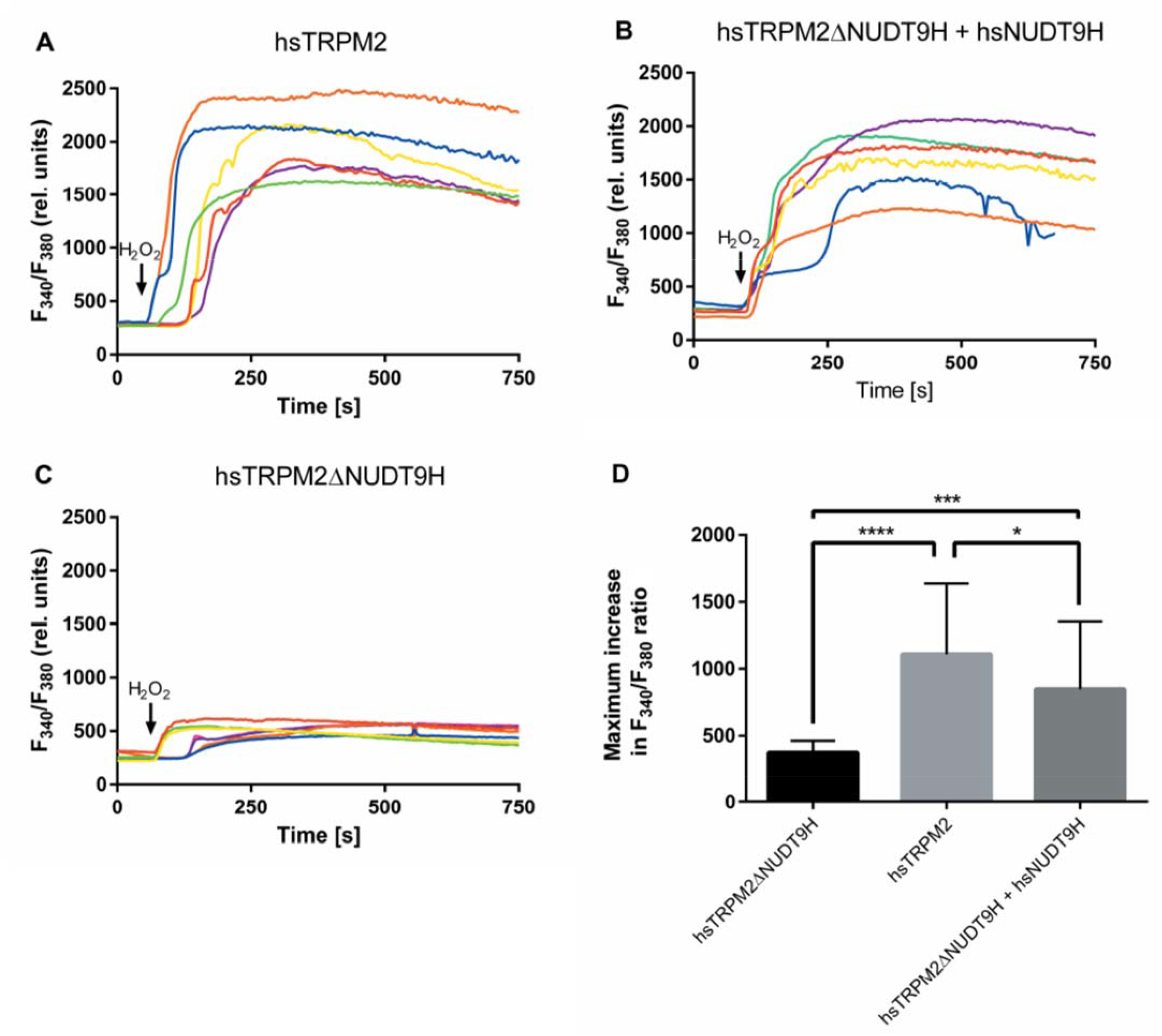
Functional analysis of in-trans interaction of hsTRPM2ΔNUDT9H and hsNUDT9H by measurements of intracellular Ca^2+^. Representative experiments of HEK-293 cells either single-transfected with wild-type hsTRPM2 as positive control (**panel A**) or co-transfected with hsTRPM2ΔNUDT9H + hsNUDT9H (**panel B**). Cells single-transfected with hsTRPM2ΔNUDT9H serve as negative control (**panel C**). Characteristic changes of [Ca^2+^]_i_ are monitored by the F_340_/F_380_ ratio over time and stimulation was performed by addition of H_2_O_2_ (10mM) to the bath solution at time points marked with arrows. Note the characteristic plateau-like increases in [Ca^2+^]_i_ after stimulation with H_2_O_2_ in panels A and B. Cells transfected with hsTRPM2ΔNUDT9H alone **(panel C)** show only a minor increase in [Ca^2+^]_i_ which corresponds to that of mock-transfected cells (not shown). In cells co-transfected with hsTRPM2ΔNUDT9H and hsNUDT9H, stimulation with H_2_O_2_ induces similar increases in [Ca^2+^]_i_ as observed with wild-type hsTRPM2, an indication for functional in-trans interaction. Maximum increases in F_340_/F_380_ ratio are summarized in **panel D**. Data are expressed as the mean ± SEM. and statistical analysis was performed with two-tailed Mann-Whitney-test (n = 12-13; ***P<0.005, ****P<0.0001, ns = not significant).

**Figure 5.**
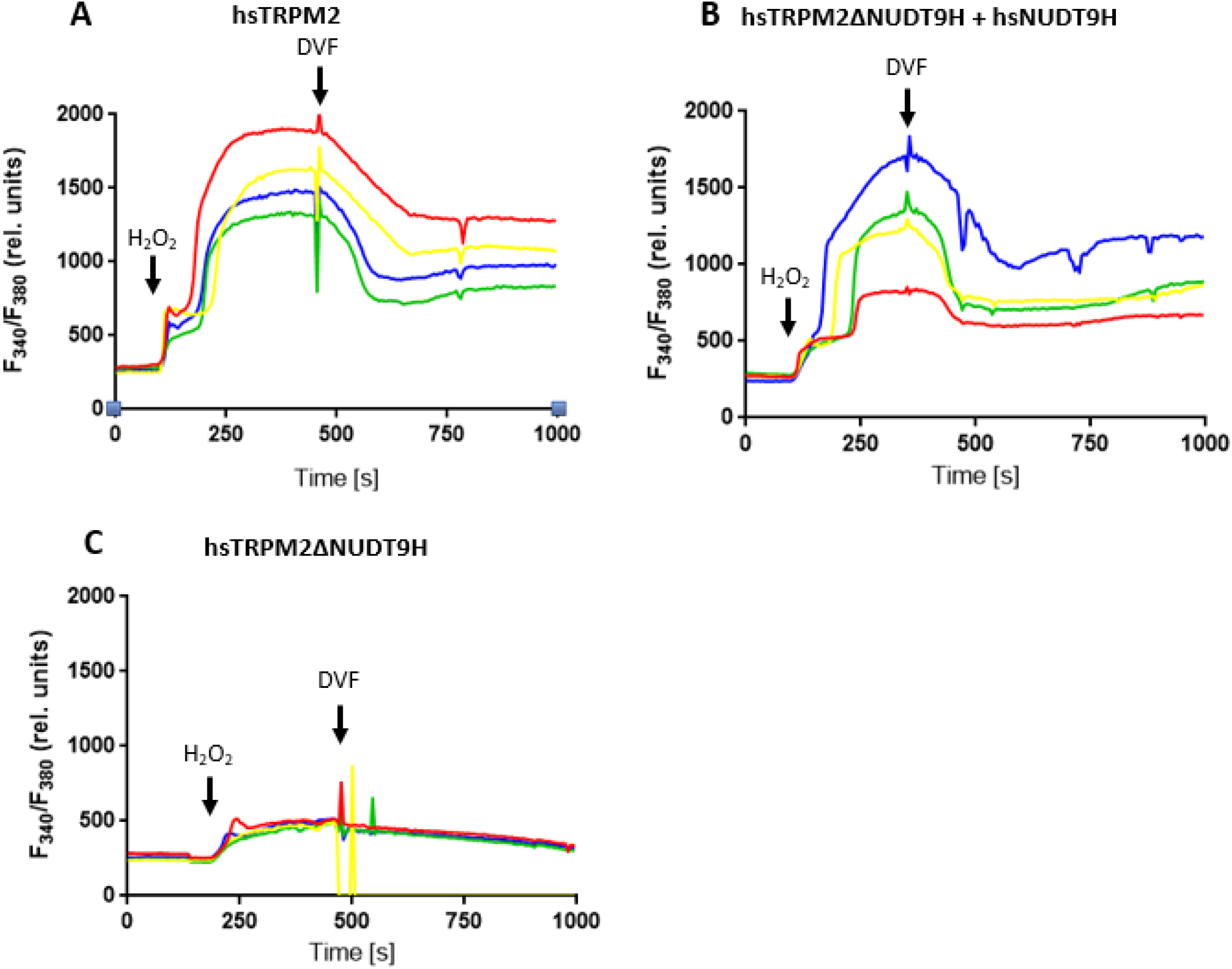
Depletion of extracellular Calcium suppresses H_2_O_2_ -induced responses of separately expressed hsTRPM2 and co-expressed hsTRPM2ΔNUDT9H + hsNUDT9H. Representative experiments of HEK-293 cells transfected as indicated (**panels A-C**). Characteristic changes of [Ca^2+^]_i_ are monitored by the F_340_/F_380_ ratio over time and stimulation was performed by addition of H_2_O_2_ (10mM) to the bath solution at time points marked with arrows. After replacing the standard bath solution (containing 1.2mM Ca^2+^) with a divalent-free bath solution (DVF; containing 10mM EGTA; indicated by arrows) the former H_2_O_2_-induced increase in [Ca^2+^]_i_ declines in cells either single-transfected with wild-type hsTRPM2 (panel A) or co-transfected with hsTRPM2ΔNUDT9H + hsNUDT9H (panel B). Cells transfected with hsTRPM2ΔNUDT9H as negative control neither respond to H_2_O_2_ nor to the replacement of the standard bath solution with divalent-free bath solution (panel C). Each of the experiments was repeated at least three times obtaining similar results.

### Functional analysis of in-trans interaction by whole-cell patch clamp analysis

For further analysis of the functional in-trans interaction between NUDT9H domain and channel domain we studied HEK-293 cells co-transfected with hsNUDT9HΔNUDT9H +hsNUDT9H with the patch-clamp technique in conventional whole-cell mode. Stimulation was performed either by ADPR (concentrations of 0.15 or 0.6 mM) applied to the cytosolic side of the plasma membrane by infusion through the patch pipette, or by H_2_O_2_ (10 mM) applied extracellularly to the standard bath solution. Representative experiments are shown in **Fig. 6**. Currents in the inward direction developed gradually during infusion of ADPR into co-transfected cells (**Fig. 6A, B)** and at a holding potential of -60 mV reached current densities (pA/pF; mean ± SEM) of 301.6 ± 70 at 0.15 mM ADPR (n = 6) and 457.5 ± 160 at 0.6 mM ADPR (n = 10; P<0.01). These currents show almost linear current voltage relation (not shown) and were reversibly blocked when extracellular Na^+^ was replaced by the large impermeable cation NMDG which suppresses the inward but not the outward component of the currents. There is a characteristic delay before reaching maximum current amplitudes which depends on the effective concentration of the agonist, as previously described for hsTRPM2 (Perraud et al., 2001). Stimulation with H_2_O_2_ induced similar currents in co-transfected cells with current densities of 198.5 ± 93 pA/pF (mean ± SEM, n = 4; P<0.01) and a latency of current onset of about 3 to 5 min. Again, inward currents were reversibly blocked by NMDG (**Fig. 6C**). The characteristic latency of current onset has been explained previously by gradual accumulation of intracellular ADPR released during oxidative-stress (H_2_O_2_) mediated apoptosis (Perraud et al., 2005). The observed current characteristics are in good agreement to previous studies on hsTRPM2 in various laboratories including our own (e. g. Perraud et al., 2001, 2005; Wehage et al., 2002; Starkus et al., 2007; Kühn et al., 2015, 2019a).

**Figure 6.**
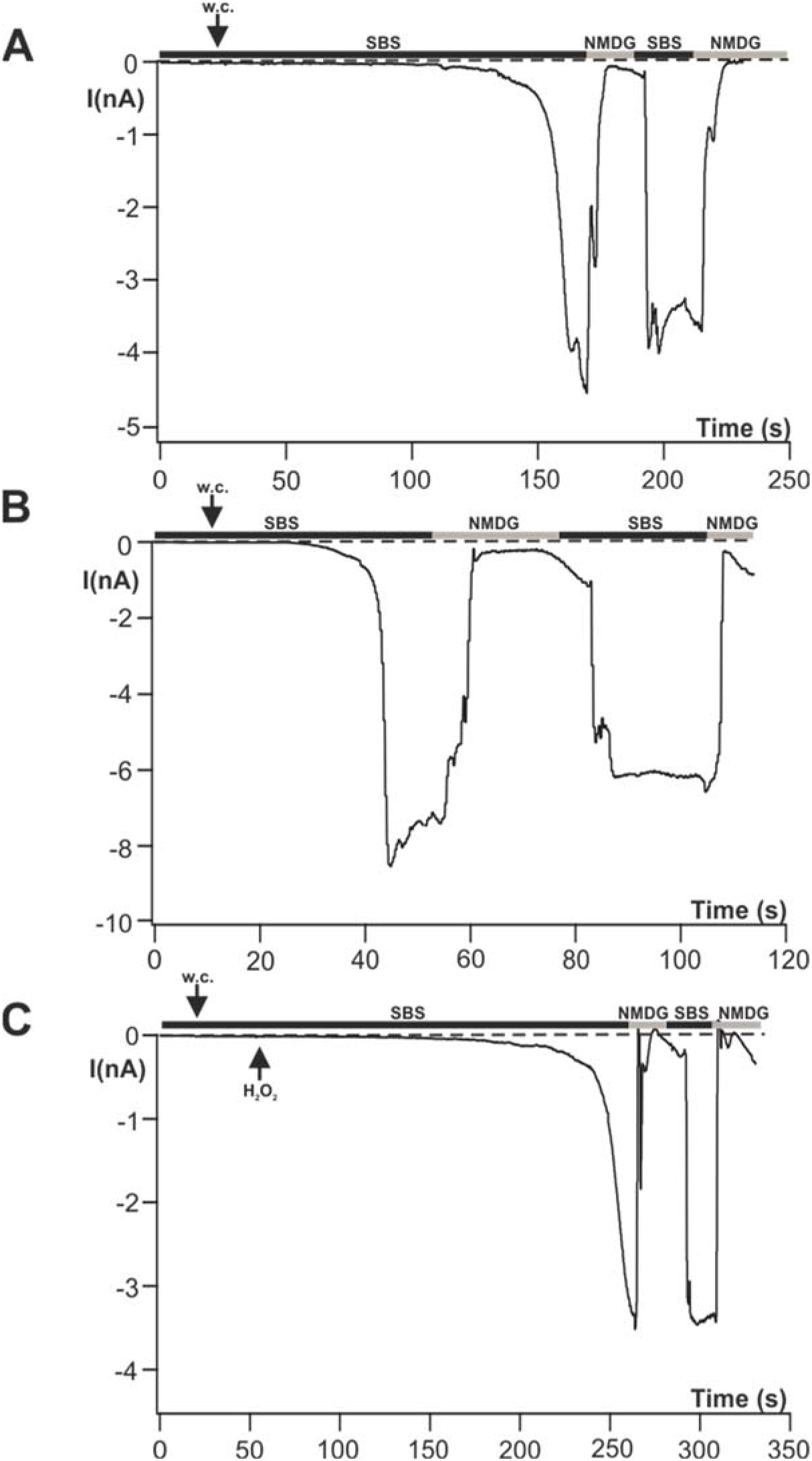
Functional analysis of in-trans interaction of hsTRPM2ΔNUDT9H and hsNUDT9H by whole-cell patch clamp recordings. Representative experiments of HEK-293 cells co-transfected with hsTRPM2ΔNUDT9H **+** hsNUDT9H. Currents were either elicited with ADPR applied through the patch-pipette in concentrations of 0.15 mM **(panel A)** and 0.6 mM **(panel B)** or with H_2_O_2_ (10 mM) dissolved in standard bath solution (SBS, **panel C**). Intracellular concentration of Ca^2+^ was adjusted to 1 µM. After establishing whole-cell configuration (ʺw.c.ʺ) a steadily increasing inward current develops, the time course of which depends on the available concentration of ADPR. Substitution of extracellular Na^+^ with the impermeable cation NMDG blocked the inward currents. Several independent experiments gave similar results (n = 10 for stimulation with 0.6 mM ADPR; n = 6 for 0.15 mM ADPR; n = 4 for H_2_O_2_).

In strong contrast, no currents were detected, after stimulation with 0.6 mM ADPR (intracellular 1 µM Ca^2+^), in HEK-293 cells expressing hsTRPM2ΔNUDT9H, the human TRPM2 channel lacking the C-terminal NUDT9H domain (**Fig. 7A**). The obtained current densities of 3.86 ± 2.9 pA/pF (mean ± SEM; n = 7) were not significantly different (P<0.005) to the Mock-control (not shown). This result is in accordance with previous data (Hara et al., 2002; Kühn et al., 2016; Huang et al., 2019). As an additional negative-control we also tested HEK-293 cells expressing hsNUDT9H, i.e. the NUDT9H domain of hsTRPM2 expressed as independent protein. As expected, no ADPR-dependent currents were generated in this case either (**Fig. 7B**).

**Figure 7.**
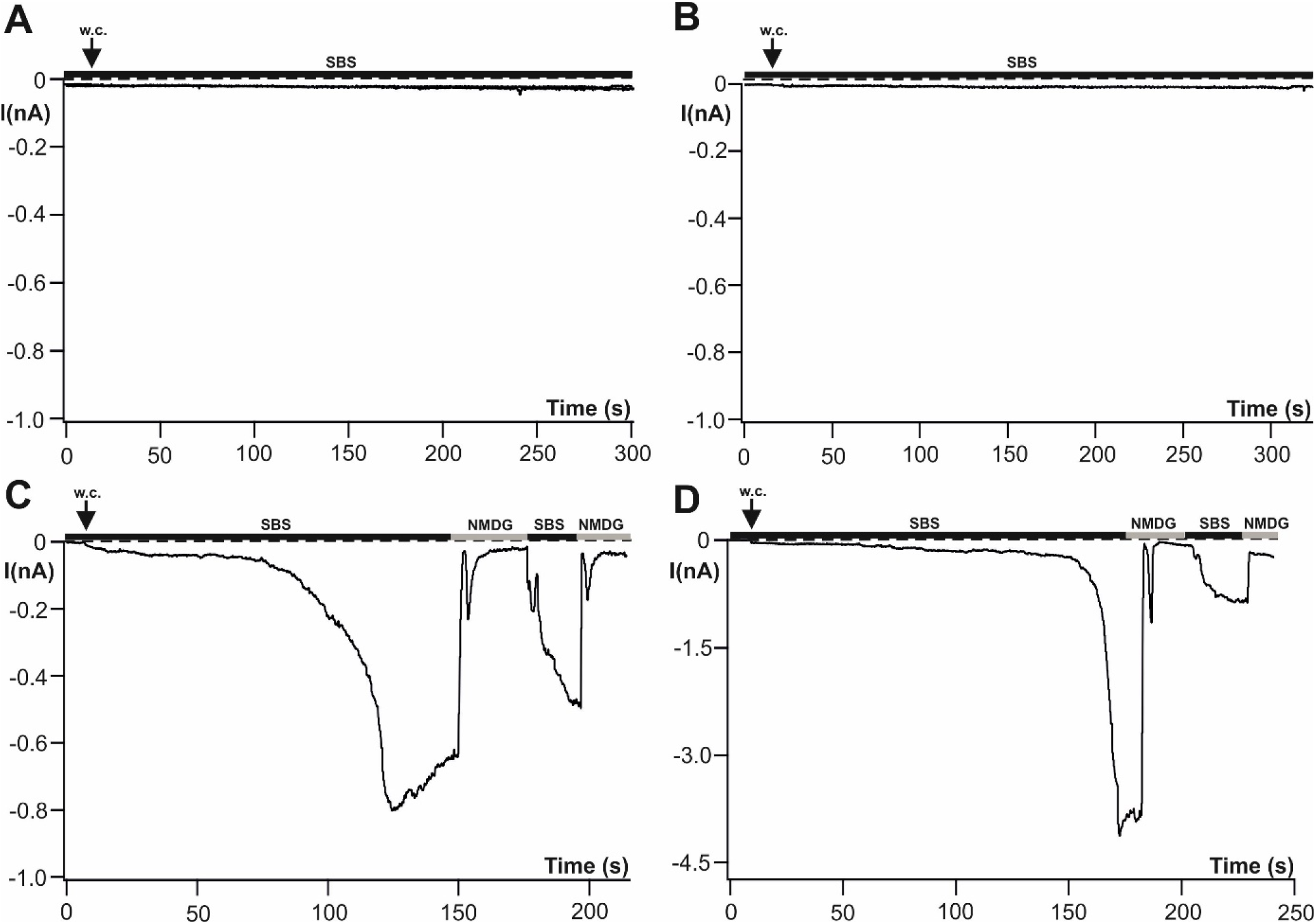
Control measurements with HEK-293 cells either expressing hsTRPM2ΔNUDT9H or hsNUDT9H alone or co-expressing hsTRPM2ΔNUDT9H + mutated hsNUDT9H. Representative whole-cell patch clamp recordings of separately expressed hsTRPM2ΔNUDT9H **(panel A)** or hsNUDT9H **(panel B)** Same protocol and abbreviations as described in Fig. 6. No current development is detectable during stimulation with ADPR (0.6 mM). Several independent experiments gave similar results (n = 7 for hsTRPM2ΔNUDT9H; n = 4 for hsNUDT9H). Examples of sporadically occurring positive responses after stimulation with ADPR (0.6 mM) in cells co-transfected with hsTRPM2ΔNUDT9H + hsNUDT9H-N1326D are depicted in **panels C and D**. In the majority of these experiments, however, no current development was detectable (statistical analysis see text).

The corresponding current densities (mean ± SEM) were 12.1 ± 11.6 pA/pF (n = 4) and not significantly different (P<0.005) to the Mock-control (not shown).

Taken together, the results obtained clearly demonstrate that the NUDT9H domain of hsTRPM2 can direct channel activity also as separately co-expressed protein.

### How does the mutation N1326D affect the in-trans interaction

Besides the deletion of the entire NUDT9H domain there are several point mutations of NUDT9H which also abolish function of the full-length TRPM2 channel (e.g. Fliegert et al., 2017; Yu et al., 2017; Huang et al., 2019). The relevant amino acid residues within the NUDT9H domain almost exclusively are involved in the binding of ADPR. However, one point mutation has been described which is located outside of these regions but still induce a loss-off-function phenotype of hsTRPM2 (Kühn & Lückhoff, 2004). The original asparagine residue at this position is strongly conserved in both invertebrate and vertebrate TRPM2 channels but is essential only for channel activity of the latter group (Kühn et al., 2016; Kühn et al., 2020). Strikingly, only the function of vertebrate TRPM2 channels essentially depends on the interaction between NUDT9H and channel domain (Kühn et al., 2016; Kühn et al., 2019b; Tóth et al., 2020; Huang et al., 2023). As already shown in our Co-IP experiments, the point mutation N1326D attenuates the binding between separately expressed NUDT9H and channel domain (**Fig. 2**). Therefore, the question arose to what extent the negative impact of N1326D on channel function is also manifested in trans. For this purpose, we performed whole-cell patch clamp analysis with HEK-293 cells co-expressing hsTRPM2ΔNUDT9H and hsNUDT9H-N1326D (**Fig. 7A, B**).

As a result, we got inconsistent results. Only 3 out of 14 cells examined showed significant ADPR-dependent current development. However, these positive currents were comparable to the currents obtained by co-expression of hsNUDT9HΔNUDT9H and wild-type hsNUDT9H (**Fig. 7A, B**). The quantification revealed current densities (mean ± SEM) of 92 ± 61 pA/pF for the positive responses (n = 3) and 5.5 ± 3.6 pA/pF for negative responses (n = 11; P<0.01). As a negative control we tested again HEK-293 cells single-transfected with hsTRPM2ΔNUDT9H and obtained current densities of 2.9 ± 0.8 pA/pF (n = 6; P<0.01).

## Discussion

The functional interaction between NUDT9H and channel domain of human TRPM2 now has been characterized by several cryo-EM analyzes (Wang et al., 2018, Huang et al., 2019, 2023). In our present study, we fully corroborate this interaction both structurally and functionally using an alternative method in a cellular context. We detect protein-protein interaction between separately expressed NUDT9H and C-terminally truncated human TRPM2 channel with two independent methods. Moreover, by co-expression of hsNUDT9H we can demonstrate the restoration of ADPR-dependent channel function of truncated hsTRPM2. This recovery of channel function also includes activation by oxidative stress, i.e. indirect activation by intracellular accumulation of ADPR, as detected by calcium imaging as well as whole-cell patch clamp measurements. The corresponding analysis of point mutation N1326D in the hsNUDT9H domain clearly shows a reduced protein-protein interaction as well as an impaired restoration of channel function. However, this result distinctly differs from the total loss-of-function phenotype induced by this mutation in full-length hsTRPM2 (Kühn & Lückhoff, 2004).

The comprehensive cryo-EM data of TRPM2 collected so far do not produce a consistent picture. For example, different interaction patterns between NUDT9H domain and channel domain have been described for the two species variants of human and zebrafish (hsTRPM2 and drTRPM2; Huang et al., 2018; Wang et al., 2018; Huang et al., 2019, 2020). However, functional studies with chimeras between hsTRPM2 and drTRPM2 show that the NUDT9H domains in principle can be swapped between the two orthologues without loss of channel function (Kühn et al., 2019b). In a recent cryo-EM study, a tetramerization of the intrinsic NUDT9H domains of archaic variants of TRPM2 is considered to be the key intermediate step for rotation of the channel subdomains MHR3/4, which ultimately induces opening of the channel pore. However, the deletion of the NUDT9H domains does not significantly affect ADPR-dependent channel activation of archaic TRPM2 (Kühn et al., 2016; Tóth et al., 2020; Huang et al., 2023). Moreover, a critical rotation of MHR3/4 was also detected in modern variants of TRPM2 (e.g. hsTRPM2), but here the preceding tetramerization step of the NUDT9H domains is absent (Huang et al., 2023). For the human NUDT9 enzyme a monomeric form of organization is assumed (Shen et al., 2003; Perraud et al., 2003; Gattkowski et al., 2021). Since the short stretch of amino acid residues critical for tetramerization of NUDT9H is absent in modern TRPM2 variants (Huang et al., 2023), it can be expected that our separately expressed hsNUDT9H domain also interacts as a monomer (Gattkowski et al., 2019). However, our results do not provide any information on how many monomers of hsNUDT9H are required to activate the tetrameric channel fragment hsTRPM2ΔNUDT9H. A structural analysis of intact aggregates of hsTRPM2dNUDT9H and hsNUDT9H could shed more light on this issue. At least, the data obtained from our Co-IP experiments indicate a stable interaction between both proteins.

Interestingly, the cryo-EM study of Huang and coworkers (2023) revealed that in archaic TRPM2 channels only the core subdomain of NUDT9H interacts with the channel domain, while in modern TRPM2 channels the cap subdomain of NUDT9H is also involved. This finding supports our current as well as previous observations, that point mutation N1326D localized within the cap subdomain impairs the function of hsTRPM2 (Kühn & Lückhoff, 2004). In contrast, the corresponding mutation N1365D does not affect channel activity in the archaic species variant from *Nematostella vectensis* (Kühn et al., 2016).

Even with the currently advanced state of knowledge, in some way the ADPR-dependent activation of TRPM2 remains a mystery. On the one hand, there are archaic species variants of TRPM2 that can exclusively be activated via the N-terminal ADPR binding pocket. On the other hand, the same N-terminal ADPR binding site is also present in modern TRPM2 variants, however, ADPR binding via the NUDT9H domain seems to be the crucial step for channel activation here. In case of archaic TRPM2 channels, the present interpretation is consistent: An ADPR-sensitive channel domain is coupled to a moderately active ADPRase, and thus manifests a negative feedback mechanism leading to a limited influx of Ca^2+^ and Mg^2+^ into the cell (Kühn et al., 2017; Huang et al., 2023). Likely, this process is reflected by the observed rapid current inactivation of nvTRPM2 along with oscillations of intracellular Ca^2+^ (Kühn et al., 2016). But how do both domains cooperate to gate human TRPM2? Based on the typical long-lasting open times of hsTRPM2, one has to assume that the NUDT9H domain produces an opposite effect as observed in archaic TRPM2 channels i.e. it facilitates the open conformation of the channel pore. While the ADPRase function is inactive (Iordanov et al, 2016), the NUDT9H domain of hsTRPM2 somehow must support channel activation via the N-terminal ADPR binding site. It is quite possible that the cap subdomain of hsNUDT9H plays an important role in this process.

One would expect that in the tetrameric hsTRPM2 channel four covalently bound NUDT9H domains differently (e.g. less flexibly) interact with the channel, if compared to one or more co-expressed NUDT9H domains. In any case, a spontaneous tetramerization of the separately expressed NUDT9H domains is very unlikely for reasons as outlined above. According to our experimental data, channel activation of wild-type hsTRPM2 via the in-trans interaction mode is almost indistinguishable from that of the in-cis interaction mode (full-length hsTRPM2). However, this is not the case for the point mutation N1326D. Here we observed a difference between the two interaction modes. Although the co-expression of the mutated NUDT9H domain in most cases abolishes channel function, significant activation occasionally occurs. This is not the case in full-length hsTRPM2, where the same point mutation induces a total loss-of-function phenotype (Kühn and Lückhoff, 2004; Du et al., 2009). This finding possibly indicates that the less stringent in-trans interaction only partially causes loss of function, while the in-cis interaction is not flexible enough to tolerate this modification.

In a previous study we obtained similarly inconsistent results regarding channel activation with the chimera hsTRPM2drNUDT9H, where the original NUDT9H domain of hsTRPM2 was exchanged by the NUDT9H domain of the species variant of zebrafish (drTRPM2; Kühn et al., 2019b). In analogy to our present interpretation, the erratic mode of activation there likewise could be explained by a suboptimal interaction, this time between channel and the foreign NUDT9H domain

In conclusion, our data indicate that the NUDT9H domain of the human TRPM2 channel can also crucially contribute to channel function as a separately expressed protein. The precise mechanism of this in-trans interaction has to be elucidated in further experiments. For example, the question arises as to whether this represents a coordinated interaction between four separate NUDT9H domains and the tetrameric channel or whether the stoichiometry is completely different. Due to the experimentally separate consideration of structural interaction and functional effects in our approach, one could also test such critical mutations, which e.g. selectively impair function without significantly disturbing protein-protein interaction. The analyzed mutation N1326D represents an example where both criteria seem to be equally negatively affected. In addition, a separate expression of orthologous NUDT9H domains could bring a further gain in knowledge. The co-expression of nvTRPM2ΔNUDT9H and separate NUDT9H domains with and without ADPRase function has already been successfully tested (Kühn et al., 2016). In any case, the main goal of future efforts should be to exactly pinpoint the mechanisms of the two different pathways of ADPR-dependent channel activation observed in archaic and modern TRPM2 channels, respectively.

## Acknowledgements

The study was supported by the Deutsche Forschungsgemeinschaft (DFG, Grant KU 2271/4-3 to FJPK). We thank Marina Wolf for expert technical assistance.

## Author contributions

FJPK conceived the study, designed and generated expression constructs, performed whole-cell patch clamp experiments and wrote the manuscript. WE carried out molecular biological experiments, site-directed mutagenesis, Co-IP experiments, proximity ligation assays, confocal microscopy and calcium imaging experiments. FJPK and WE conducted data analysis and prepared figures.

## Data availability

The datasets generated during and/or analysed during the current study are available from the corresponding author on reasonable request.

**Notes:** The authors declare no competing financial interests.

